# Skin microbiome attributes associate with biophysical skin aging

**DOI:** 10.1101/2023.01.30.526239

**Authors:** Wei Zhou, Elizabeth Fleming, Guylaine Legendre, Lauriane Roux, Julie Latreille, Gaëlle Gendronneau, Sandra Forestier, Julia Oh

## Abstract

Two major arms of skin aging are changes in the skin’s biophysical conditions and alterations in the skin microbiome. This work partitioned both arms to study their interaction in detail. Leveraging the resolution provided by shotgun metagenomics, we explored how skin microbial species, strains, and gene content interact with the biophysical traits of the skin during aging. With a dataset well-controlled for confounding factors, we found that skin biophysical traits, especially the collagen diffusion coefficient, are associated with the composition and the functional potential of the skin microbiome, including the abundance of bacterial strains found in nosocomial infections and the abundance of antibiotic resistance genes. Our findings reveal important associations between skin biophysical features and aging-related changes in the skin microbiome and generate testable hypotheses for the mechanisms of such associations.

## Introduction

Human skin has multiple environmental niches compactly assembled on a surface area of 1-2 square meters and is home to millions of microbes. Mechanistically, skin microbes have long been known to contribute to skin physiology, including training and modulating the skin immune system, influencing the surface pH of the skin through lipid metabolism, and providing colonization resistance against pathogens ^1–3^. Recently it has also been shown that the skin microbiome can regulate the formation, repair, and function of the epidermal barrier ^4–6^.

Conversely, numerous behavioral and host intrinsic factors shape the skin microbiome composition, including the substantial physiologic changes inherent to skin aging. Our initial metagenomic characterizations of older adult cohorts ^7^suggest that the skin microbiome varies substantially in the long term, especially during the process of aging. Skin aging results from a multi-factorial process that includes extrinsic (engendered by external stressors such as UV and pollution exposure, for example) and intrinsic (genetic clock, hormonal changes, etc.) factors ^8,9^. Skin aging is characterized by alterations in the tissue structure, including epidermal thinning, a flattening of the junction between the epidermis and dermis, and the loss of elastin and collagen ^8,9^. These tissue changes are subsequently reflected biophysically as alterations in parameters such as skin capacitance or collagen quality and quantity measurements, as well as clinical parameters such as wrinkles, sagging, or color irregularities^10^.

While these physiologic changes undoubtedly result in corresponding changes in skin microbiome composition ^11–17^, a well-controlled, high-resolution characterization taken together with detailed biophysical skin measurements is lacking. On one hand, confounding factors such as sex, ethnicity, geography, and health condition can all affect skin microbiome^18^, which increase data dimensionality and decrease statistical power ^7^. On the other hand, while previous investigations revealed aging-related skin microbiome changes at phylum-to genus-resolution ^11,14–18^, studies by our group and others have shown that the aging-related diversity of the skin microbiome is often manifested at species-, strain-, or gene-level ^7^. Moreover, such high-resolution diversity has potential health implications. For example, we found that a phylogenetic clade of *Staphylococcus (S*.*) epidermidis*, which contains strains associated with nosocomial infections, was significantly enriched in adults living in skilled nursing facilities, compared to younger adults ^7^. We hypothesize that these high-resolution changes in the skin microbiome are not only a biomarker of aging but also associated with aging-induced biophysical changes in the skin.

Thus, we used shotgun metagenomic sequencing to characterize species, strain, and gene content distribution of the skin microbiome in older vs. younger adults. We also collected host attributes – BMI and biophysical parameters, including dermis water content (DWC), skin capacitance (SC, a measurement of stratum corneum water content), and collagen diffusion coefficient (CDC, a measurement of collagen quality and quantity). To reduce data dimensionality and control for confounding factors of the skin microbiome composition, we confine the investigation to only the facial skin microbiome of healthy Caucasian women in Paris area. Using this well-controlled dataset, we found significantly different facial microbiome structures in younger and older adults. More importantly, we found that much of this difference can be attributed to the variation in biophysical parameters, especially CDC.

## Results

### Changes in biophysical characteristics during aging

We recruited 26 younger women (20-26 yo) and 25 older women (54-60 yo). We collected microbiome samples via swabs from both cheeks (except for one older woman, V2G), hygiene survey, body mass index (BMI), and biophysical parameters including collagen diffusion coefficient (CDC), skin capacitance (SC), and dermis water content (DWC) (Supplementary Table S1). Aging-related structural alterations of the skin resulted in measurable changes in biophysical parameters. CDC was significantly higher in the younger subjects than the older subjects (Figure 1A, Wilcoxon test p=5.7×10^−5^), while BMI was significantly higher in the older subjects (Figure 1A, Wilcoxon test p=0.011). SC and DWC were not significantly different between the two groups (Figure 1A). Among the host attributes, CDC exhibited a significant and consistent correlation with SC in both younger (Pearson’s *r*=0.59, q=1.26×10^−5^) and older (Pearson’s r=0.41, q=0.0095) subjects (Figure 1B). Significant correlations were also observed for CDC with BMI (Pearson’s *r*=-0.44, q=0.0021) and DWC (Pearson’s *r*=0.33, q=0.028), but only within the younger women. However, the variance inflation factors of the host attributes were low (Supplementary Table S1), indicating that multicollinearity should not significantly influence downstream statistical analyses.

**Figure 1.**
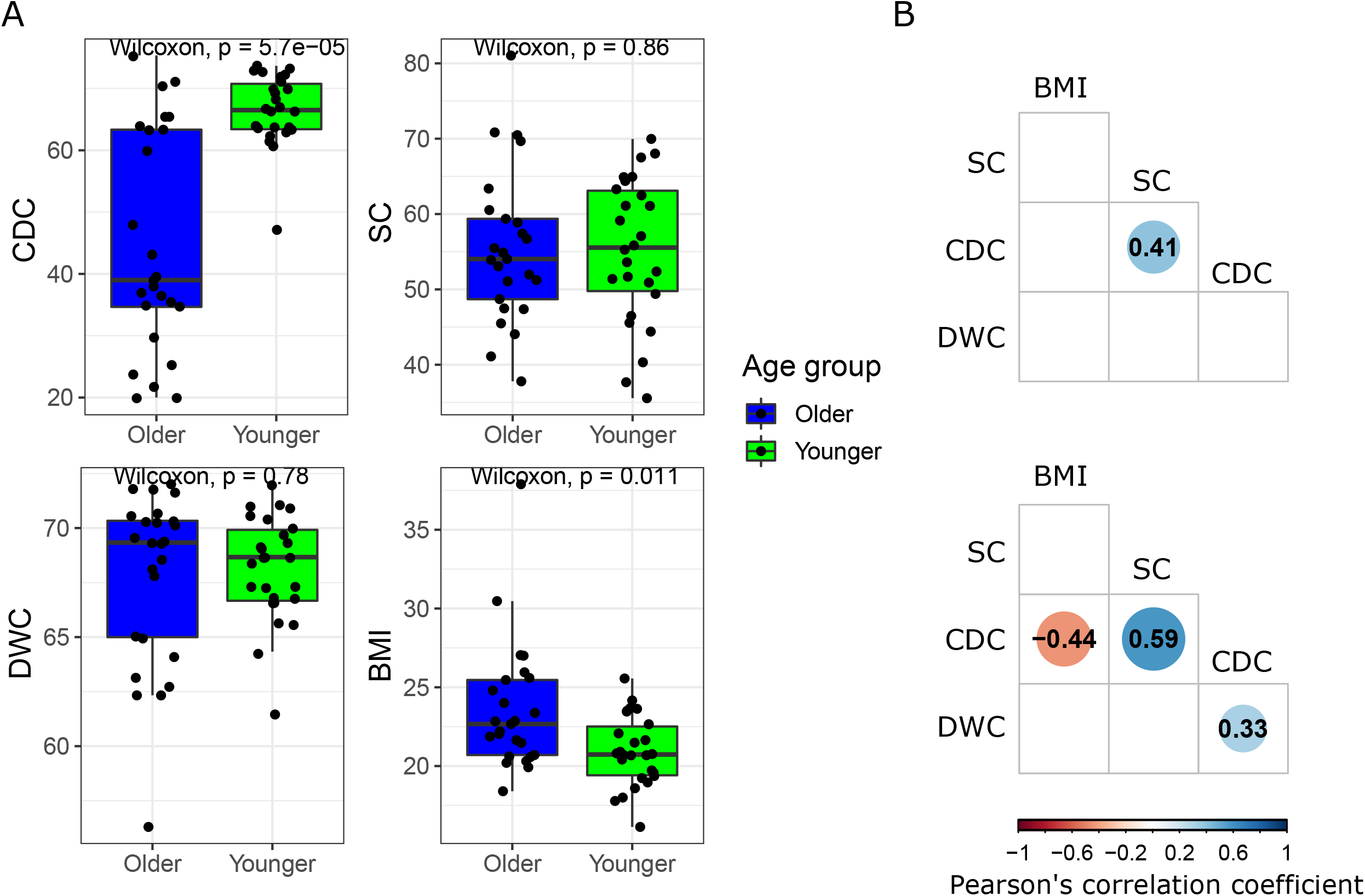
Skin biophysical traits of the older and younger subjects. A, Comparison of the biophysical traits between age groups. B, Correlation between the biophysical traits. Only statistically significant correlations were shown (q<0.1). The sizes and colors of the circles were proportional to Pearson’s correlation coefficient (the number inside the circle).

### Changes in species-level community structure during skin aging

Extraction and shotgun sequencing of the facial microbiome swabs resulted in, after preprocessing, 7.4±6.6 million quality-filtered paired-end microbial reads per sample (Supplementary Table S2, maximum: 29.6 million, minimum: 0.73 million, median: 5.0 million). *Cutibacterium (C*.*) acnes* (79.4±17.4%) was the dominant species in all subjects, followed by *S. epidermidis* (4.9±6.5%) and *Corynebacterium (Cor*.*) kroppenstedtii* (2.5±5.7%) (Figure 2A), consistent with previously observed microbial community structure at sebaceous sites ^19,20^. Younger women had a significantly higher proportion of *C. acnes* than older women (84.9±15.9% in younger subjects vs 73.7±17.3% in older subjects, Wilcoxon test p=0.0054), as well as moderately lower proportions of *S. epidermidis* (4.1±7.1% in younger subjects vs 5.8±5.8% in older subjects, Wilcoxon test p=0.11) and *Cor. Kroppenstedtii* (1.3±5.0% in younger subjects vs 3.7±6.2% in older subjects, Wilcoxon test p=0.060), concordant with previous studies from our group and others ^7,15,18^.

**Figure 2.**
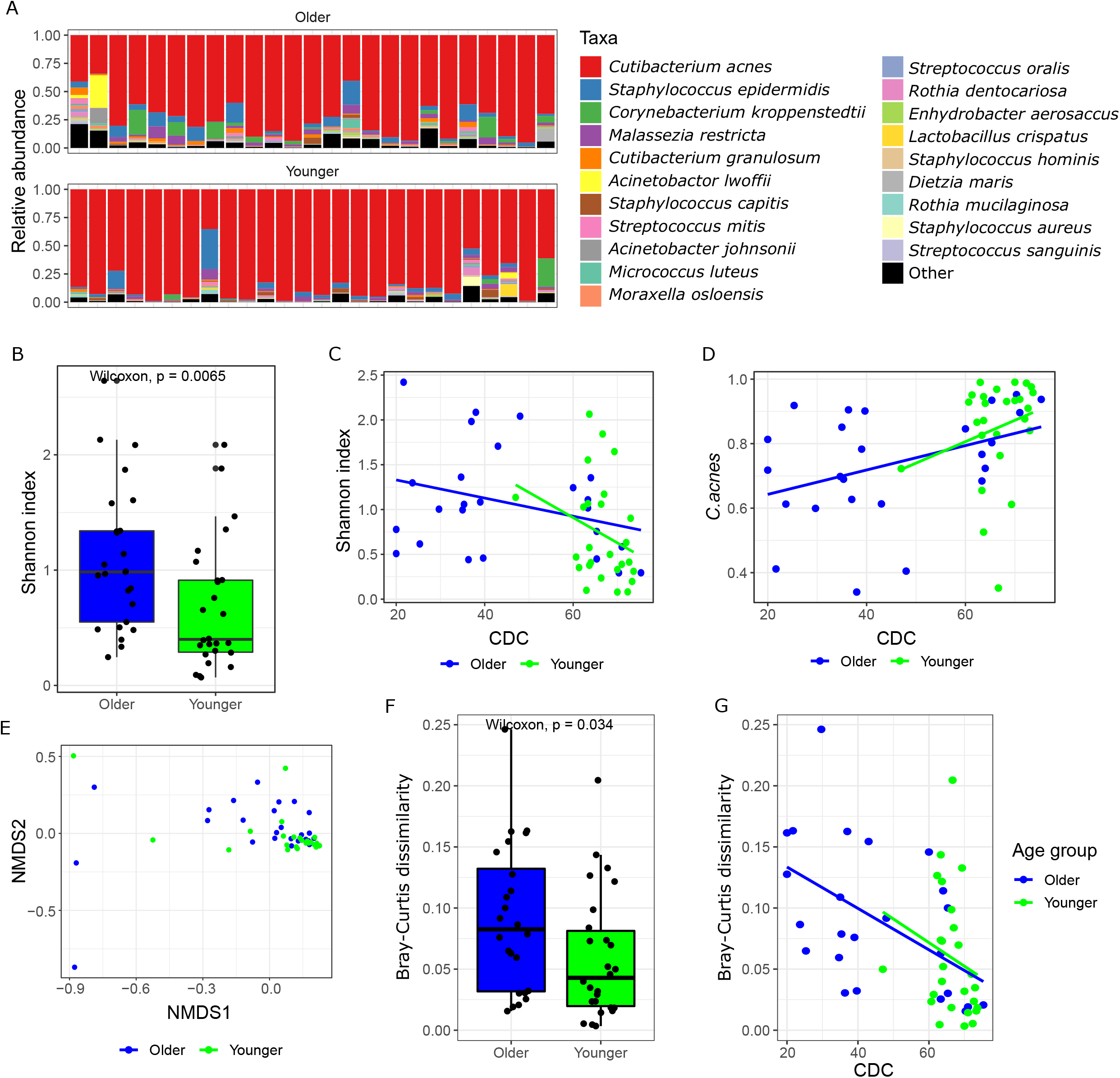
Diversity of microbial species in the older and younger skin microbiome. A, Distribution of the top twenty most abundant microbial species. B, Species-level alpha diversity of the younger and older samples as measured by Shannon’s index. C, Association between CDC and Shannon’s index within age groups. D, Association between CDC and the relative abundance of *C. acnes* within age groups. E, non-metric multidimensional scaling (NMDS) plot of the younger and older samples at species resolution. F, intra-personal heterogeneity in the facial microbiome between age groups. G, Association between CDC and intra-personal heterogeneity within age groups.

At the species-level, samples from younger subjects showed lower alpha diversity compared to the older subjects (Figure 2B, Wilcoxon test p<0.0065 for Shannon’s index), which is in concordance with previous studies^7,11,15,17,18^. This is at least partially and mathematically due to the dominance of *C. acnes* in the younger skin that deflates species evenness. We then investigated if the observed diversity difference between age groups can be attributed to skin biophysical parameters or BMI. To do this we fit the data to a linear model and questioned if each host attribute (i.e. age group, BMI, CDC, SC, and DWC) had a significant influence on alpha diversity when adjusted for all other variables. We found that CDC had the most significant influence on alpha diversity (ANOVA type II test p=0.085) among the variables tested (ANOVA type II test p>0.3 for all other variables), that is, the CDC difference is negatively associated with species diversity both between and within age groups (Figure 2C). Consistently, CDC is positively associated with *C. acnes* abundance (Figure 2D, ANOVA type II test p=0.086; p>0.3 for all other variables, including age group), which deflates species evenness. These findings propose a mechanistic link between aging and skin microbiome: aging decreases the production of collagen, which could result in a decrease in *C. acnes* abundance and consequently an increase in species diversity.

The two age groups have significantly different facial microbiome compositions (Figure 2E, Distance-based redundancy analysis (dbRDA) on Bray-Curtis dissimilarity p=0.0023). Similar to alpha diversity, when we investigated the influence of BMI and biophysical parameters on the facial microbiome composition, we found that only CDC has a significant effect (ANOVA type II test p=0.043; p>0.3 for all other variables, including age group). Our group previously reported an increased microbial diversity between body sites observed in subjects of older age and greater frailty^7^, here we validate the finding in the present dataset which is better controlled for confounders. We asked if the intra-personal heterogeneity (i.e. Bray-Curtis dissimilarity between microbiomes of the left and the right cheek) also differs between age groups. Notably, this intra-personal heterogeneity differs from our previous report in that it is controlled for differences in different body niches (e.g., sebaceous facial environment compared to moist foot environment). We found that the older women showed significantly higher intra-personal microbiome heterogeneity (Figure 2F, Wilcoxon test p=0.034), which is independent of aging-induced diversifications of the environmental conditions at different body sites. By breaking down the contribution of each host attribute to intra-personal heterogeneity, we found that, again, the difference between age groups can be attributed to differences in CDC (Figure 2G, ANOVA type II test p=0.016; p>0.5 for all other variables, including age group). These findings underlie the relevance of CDC in explaining microbial community structures both between and within age groups, suggesting that skin physical condition can potentially influence (or be influenced by) the assemblage of skin microbial species, a hypothesis now explored in vitro^21^.

### Changes in strain-level population structure during skin aging

Given that strain variation of keystone skin microbes has been linked with a diversity of functional phenotypes, including virulence, we examined if skin aging was associated with changes at the strain-level. We inferred the abundances of different phylogenetic clades of *C. acnes* and *S. epidermidis* in the dataset with a reference-based method previously described in Larson et al^7^. Like in our previous study, we focused on *C. acnes* and *S. epidermidis* because of their ubiquity and biological relevance in the skin. We leveraged a curated set of phylogenetically diverse reference genomes and identified 10 and 13 phylogenetic clades for *C. acnes* and *S. epidermidis*, respectively^7^ (Supplementary Figure S3). Metagenomic reads were then assigned to the most likely clade of origin using Pathoscope2, providing an estimate of the relative abundances of the phylogenetic clades in their respective population (Figure 3A).

**Figure 3.**
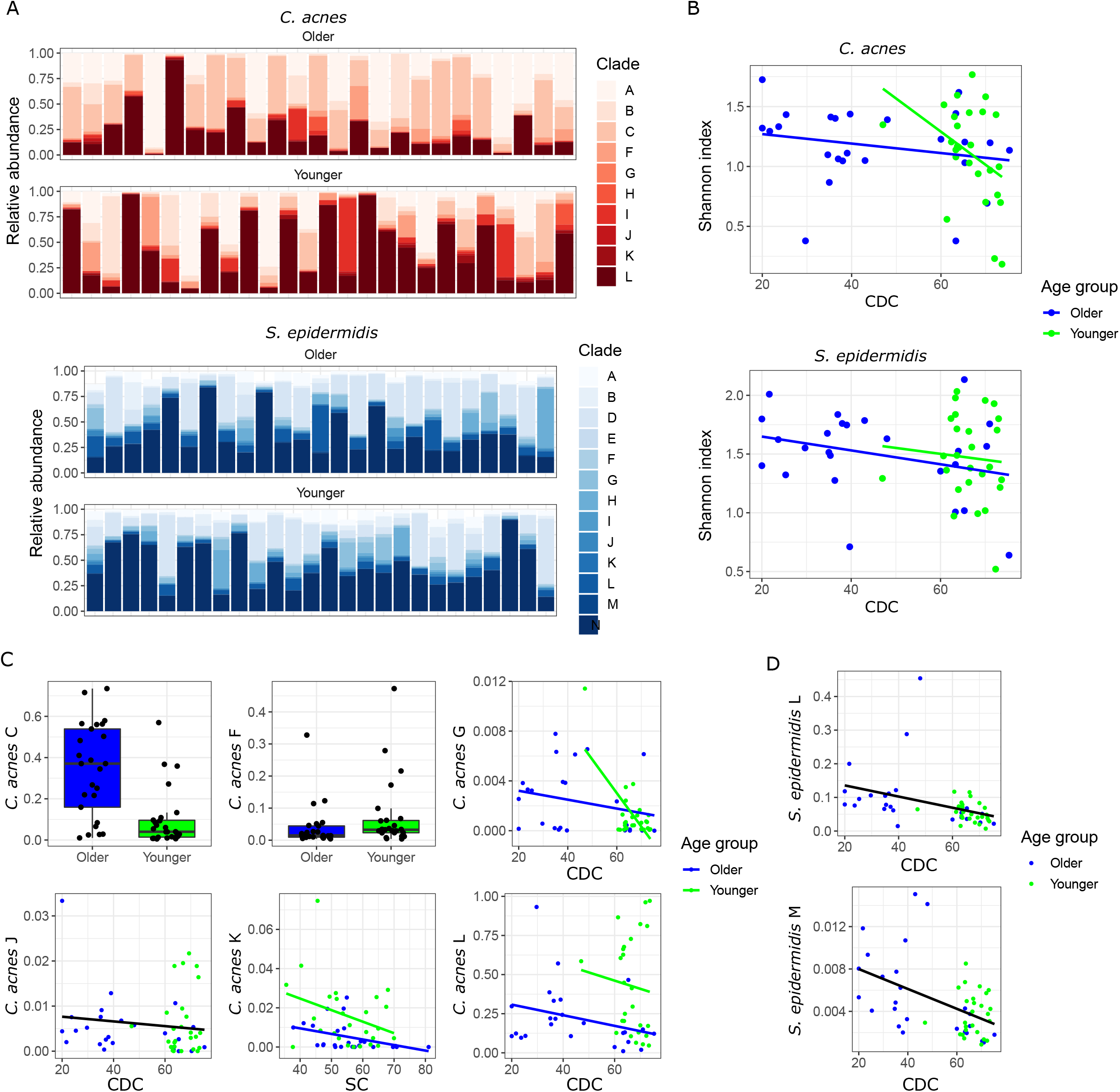
Population structuring of *C. acnes* and *S. epidermidis*. A, Relative abundances of the phylogenetic clades of *C. acnes* and *S*.*epidermidis*. B, Associations between CDC and the population diversities (measured by Shannon’s index) of *C. acnes* and *S. epidermidis* within age groups. C, Associations between *C. acnes* phylogenetic clades and host attributes. Only associations with q<0.25 were shown. D, Associations between *S*.*epidermidis* phylogenetic clades and host attributes. Only associations with q<0.25 were shown.

Neither the *C. acnes* nor the *S. epidermidis* population showed differential alpha diversity between age groups. Also, alpha diversities of the two species were not significantly correlated with any biophysical parameters measured, although CDC appeared to have a moderate effect (Figure 3B, ANOVA type II test p=0.15 and 0.10 for *S. epidermidis* and *C. acnes* respectively). We next dissected the population composition to assess individual associations between host attributes and the abundance of each phylogenetic clade using MaAsLin2^22^. Due to the high data dimensionality, we report associations with q values (Benjamini-Hochberg-controlled) lower than 0.25 – the default threshold adopted by MaAsLin2. *C. acnes* clades G (p=0.0022, q=0.054), J (p=0.018, q=0.16), and L (p=0.020, q=0.16) were negatively associated with CDC, while clade K was negatively associated with SC (p=0.036, q=0.22) (Figure 3C). In addition, Clades C (p=0.018, q=0.16), F (p=0.043, q=0.24), G (p=0.022, q=0.16), K (p=0.0051, q=0.086), and L (p=0.0012, q=0.054) also exhibited an association with age group after adjusted for other host attributes (Figure 3C), suggesting the presence of age-related factors that shaped the *C. acnes* population composition but were not measured in this study. Among the clades, only Clade C was observed to be enriched in the older women (Figure 3C, Wilcoxon test p=7.8×10^−5^, when tested between age groups while pooling all other host attributes). Clade C consisted of genomes belonging to phylotype IA1 clonal complex 4 (CC4), which was found to be enriched in patients with acne and capable of inducing inflammation and tissue damage^23–25^. On the contrary, CDC was the only variable associated with the abundance of any *S. epidermidis* clade (L, p=0.0066, q=0.21; M, p=0.001, q=0.068) at q=0.25 (Figure 3D). *S. epidermidis* clade L contains strains found in nosocomial infections and was previously shown to be enriched in older adults living in skilled nursing facilities^7^. Thus, the negative association between CDC and *S. epidermidis* clade L suggests a possible link between the falling collagen quantity and quality in older adults and the susceptibility to nosocomial infections.

### Changes in microbial functions during skin aging

Older individuals have inherently experienced greater cumulative exposure to extrinsic stressors and taken together with aging-specific intrinsic factors, we anticipate potential functional differences accumulated by the skin microbiome. We observed three differentially abundant functional pathways between the age groups at q<0.25: “phosphopantothenate biosynthesis I” (BioCyc ID: PANTO-PWY, Wilcoxon p=0.00074, q=0.22), “superpathway of sulfur amino acid biosynthesis” (BioCyc ID: PWY-821, Wilcoxon p=0.00020, q=0.059), and “superpathway of glyoxylate bypass and TCA” (BioCyc ID: TCA-GLYOX-BYPASS, Wilcoxon p=0.00063, q=0.19, and Wilcoxon p=0.0019 for GLYOX-BYPASS alone) (Figure 4A). All three pathways were enriched in samples from the older women. In particular, the glyoxylate bypass mechanism was known to be upregulated under conditions of oxidative stress^26^, consistent with the fact that elevated oxidative stress is a hallmark of skin aging^27^. When adjusted for host attributes, “Rubisco shunt” (BioCyc ID: PWY-5723, p=0.00012, q=0.094) and “superpathway of beta-D-glucuronide and D-glucuronate degradation” (BioCyc ID: GLUCUROCAT-PWY, p=0.00042, q=0.23) were found to be differentially abundant between age groups; both of which were enriched in samples from the older women (Figure 4B), representing aging-related microbiome functional changes that are not explained by the host attributes measured in this study. In addition, “superpathway of pyrimidine ribonucleosides salvage” (BioCyc ID: PWY-7196, p=4.4×10^−5^, q=0.07) was found to be associated with DWC, but the association was likely driven by one outlier (subject V48; p=0.0098 and q=0.79 when excluded).

**Figure 4.**
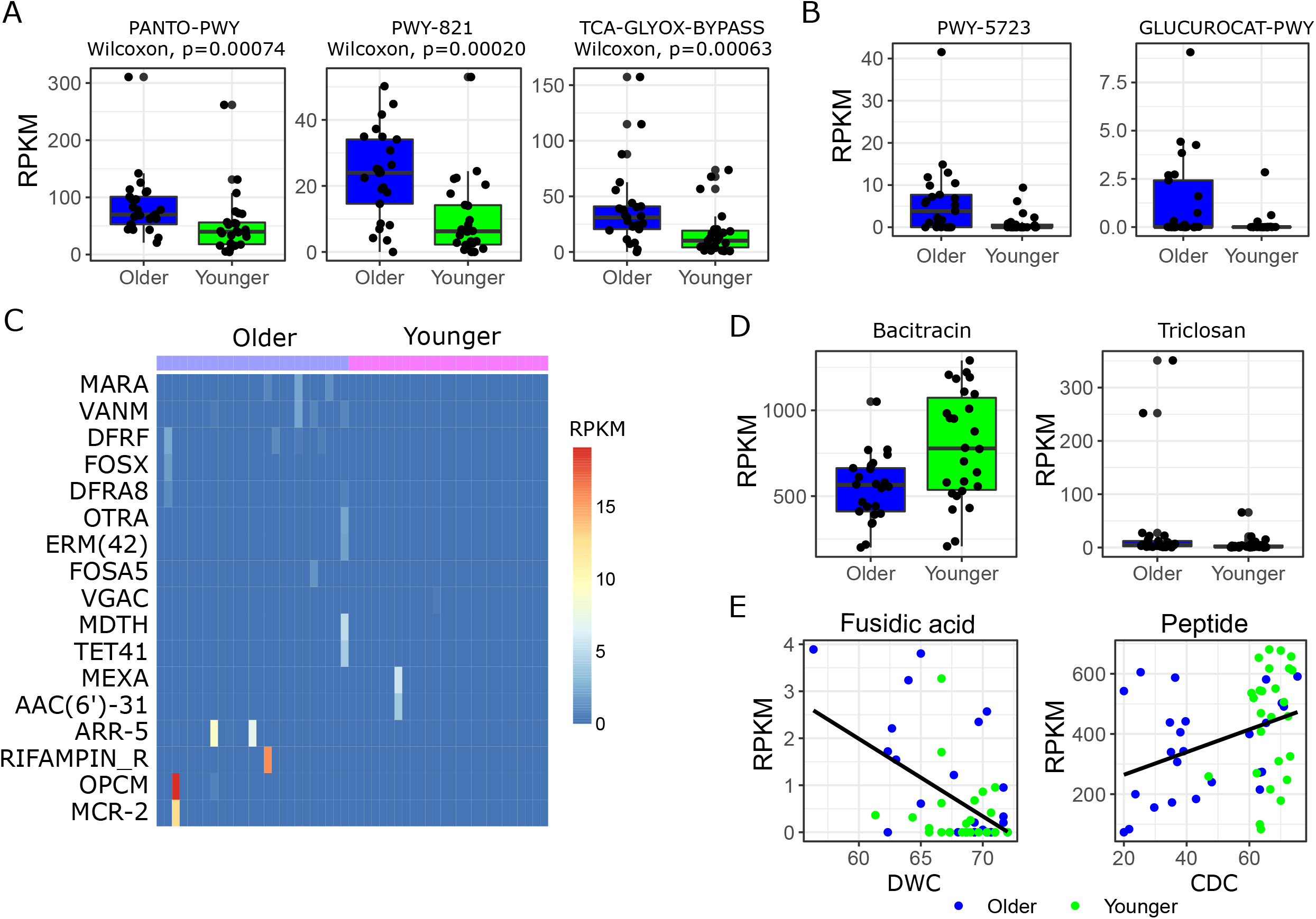
Functional diversity in the older and younger skin microbiome. A, Microbial pathways differentially abundant between age groups (q<0.25, not adjusted for host attributes). Microbial pathways were represented by their BioCyc ID. B, Microbial pathways differentially abundant between age groups (q<0.25, after adjusted for host attributes). Microbial pathways were represented by their BioCyc ID. C, Distribution of age-group-specific AR genes. RPKM: reads per kilobase of gene sequence per million reads sequenced. D, AR genes differentially abundant between age groups (q<0.25, not adjusted for host attributes). E, Associations between AR gene abundances and host attributes. Only associations with q<0.25 were shown. RPKM: reads per kilobase of gene sequence per million reads sequenced.

A particular example of clinical relevance is the accumulation of antibiotic resistance (AR) genes via cumulative exposure – topically or systemically – to antibiotics. AR is an increasing concern in treating skin infections and is a function of exposure to antibiotics. Interestingly, the overall abundance of AR in the facial microbiome of the older women was not significantly higher than the younger women (reads per kilobase of gene sequence per million reads sequenced, RPKM, 11129±3926 for the older women, and 12603±4226 for the younger women), although the former had more opportunities of exposure to antimicrobials over a lifespan. Nonetheless, out of 313 antibiotic resistance genes (AR genes) predicted from our gene catalog, 14 were specific to older subjects (i.e., present in at least one older subject while present in no younger subjects), while only 3 were specific to younger subjects (Figure 4C). Among these age-group-specific AR genes, 9 were shared by at least two older subjects, while none was shared by any two younger subjects (Figure 4C). These results revealed a richer but not more abundant AR gene reservoir in the facial microbiome of older women. In terms of differential abundance, bacitracin resistance genes were observed to be elevated in the younger women (Figure 4D, Wilcoxon test p=0.0054, q=0.14) and triclosan resistance genes in the older (Figure 4D, Wilcoxon test p=0.0054, q=0.14). When adjusted for host attributes, bacitracin resistance genes still appear enriched in the younger samples (p=0.0033, q=0.19), indicating the influence of unmeasured age-related variables. AR gene abundance is also associated with Biophysical parameters: DWC was found to be negatively correlated with the abundance of fusidic acid resistance genes (Figure 4E, p=0.0016, q=0.19; and p=0.0072, q=0.23 after removing the apparent outlier V48), and CDC was found to be positively correlated with the abundance of genes conferring resistance to antimicrobial peptides (Figure 4E, p=0.0045, q=0.19), which, like CDC, may diminish during aging^28^. Our analyses also reported BMI to be associated with AR genes towards MLS, polyamine peptide, tetracycline, and fusaric acid, but these associations were likely driven by one outlier sample (subject V9; no associations reported at q<0.25 after removing the sample).

## Discussion

Taken together, our study extended previous investigations of aging-related skin microbiome changes into species-, strain-level, and gene-level. We moreover identified new associations with biophysical parameters, especially CDC, on multiple resolutions, which are typically lacking in skin microbiome studies despite the skin microbiome’s close link to skin physiological differences. Overall, we found that skin microbiome diversity increased with age at the species level, consistent with other studies conducted on Caucasians ^11^ and other ethnicities^14–16^.

With the increased resolution provided by shotgun metagenomics, we were able to confirm our previous findings^7^ that *Cor. kroppenstedtii* was likely responsible for the observed aging-related increase in *Corynebacterium* in both Kim et al^14^ and Shibagaki et al^15^, and *C. acnes* was likely responsible for the aging-related depletion of *Cutibacterium* in Shibagaki et al^15^. Consistent with our previous finding^7^, we again presented evidence indicating that closely related strains of the same species could interact differently with the skin aging process.

However, with the additional measurement of biophysical parameters, in this study, we were able to link species and strain abundance with numerical host attributes. Although SC and DWC were not significantly different between younger and older groups, collagen quality and quantity (measured through CDC) were significantly decreased with skin aging due to its progressive degradation and decrease of production with both chronological and photo-aging^29^. Through high-resolution profiling of the skin microbiome, we found that both community structuring and strain population structuring were associated with biophysical parameters of the skin habitat. Most strikingly, CDC was able to statistically explain multiple age-related patterns observed in the microbiome. For example, CDC was associated with community diversity and structure, consistent with previous findings that skin physical properties can affect microbial distribution^30^. Moreover, we found a correlation between CDC and two phylogenetic clades of *S. epidermidis*, one of which (clade L) contained strains involved in nosocomial infections and was previously found to be enriched in adults living in skilled nursing facilities^7^. This finding converts a statistical observation into a testable hypothesis and could guide potential prevention practices to control *S. epidermidis* nosocomial infections. Contrarily, *C. acnes* strains appear to be correlated with not only the biophysical parameters but also age-related factors that were not measured in this study. As *C. acnes* is lipophilic, one variable that could affect *C. acnes* population is skin sebum level, which decreases with age, especially in women^31^. Another complication was that *C. acnes* population diversity was maintained through spatial segregation and bottlenecking across skin pores^32^ – a largely stochastic process. Despite that this mechanism could obscure non-stochastic patterns associated with host attributes, we were able to identify a significant difference in clade C abundance between age groups, thanks to the rigidly controlled dataset. *C. acnes* strains in clade C belong to phylotype IA1 and clonal complex CC4, which was pro-inflammatory and enriched in acne patients ^23–25^. As inflammation and the resulting tissue damage are considered driving forces of skin aging^33^, it can be hypothesized that *C. acnes* clade C is functionally related to skin aging. Future studies, for example *in vitro* young vs. aged skin equivalent models inoculated with young vs. aged skin microbiota (in particular, the older adult-specific strain of *S. epidermidis* and *C. acnes*) as determined in this study, would be of interest to determine if skin biophysical parameters influence or are influenced by skin microbial diversity.

Among the host attributes analyzed in this study, CDC appears to be the most profound statistically, as it can not only explain many differences between the age groups but also differences within the age groups. Therefore, we can hypothesize that CDC, as a parsimonious predictor, can functionally facilitate (or is facilitated by) age-related changes in the facial microbiome. Kim et al^21^ showed that younger Korean women carried a higher amount of skin Streptococcus compared to older Korean women. These Streptococcus species, specifically *Streptococcus (St*.*) pneumoniae* and *St. infantis*, can secrete substances that up-regulate collagen synthesis. However, Streptococcus abundance was not higher in the younger adults recruited in our study, nor was it positively correlated with CDC (data not shown). This discrepancy could be due to the differences in study population (Korean vs European Caucasian), sequencing method and resolution (16S rRNA sequencing vs shotgun metagenomic sequencing), and the difference in the age range of the recruited older women (40-53 yo vs 54-60 yo). Therefore our results suggested that the link between skin collagen and skin microbiome extends beyond the function of one genus.

Multiple metabolic pathways were enriched in the aging facial microbiome. Especially of interest is the enrichment of glyoxylate bypass pathway, as its up-regulation is observed during oxidative stress – a characteristic of skin aging. This observation is consistent with a previous study on the skin microbiome of healthy Chinese women^12^, exhibiting an age-related functional change that is shared by at least two drastically different populations. In terms of clinical relevance, we expected to observe a more diverse AR gene reservoir in the older skin microbiome, as observed previously. Indeed, the older skin microbiome did exhibit a richer collection of AR genes compared to the younger skin microbiome, but the overall abundance was not higher. One example of AR enriched in the younger skin microbiome is the resistance to Bacitracin, which is commonly applied topically to prevent minor infections. Another example is the resistance to antimicrobial peptides. Although antimicrobial peptide resistance was not statistically significantly enriched in the younger women per se, it was positively correlated with CDC, therefore indirectly associated with the aging process. Interestingly, skin collagen secretion and antimicrobial peptide production can both be traced back to functions of the dermal fibroblasts, and both diminish during aging^28^. Thus, our finding proposed a pleiotropy hypothesis: decreased dermal fibroblast activity loosens the selection pressure imposed by antimicrobial peptides and at the same time decreases collagen synthesis.

The object of our study is to address not only the effect of aging but also what aging-related host attributes, especially skin biophysical parameters, interact with the skin microbiome. Although we successfully identified CDC as an important indicator of high-resolution microbiome patterns, many microbial variables differed between age groups after adjusting for host attributes. Therefore, aging, as a pooling of biological effects, must contain other sources of variation that were not measured in this study. For example, skin sebum level significantly correlates with local community composition^34^ and can be quantified using the Sebumeter^35^; skin surface antimicrobial peptides directly shape microbial taxa frequencies and a recent study showed that skin surface antimicrobial peptides can be measured noninvasively using a skin patch test^36^. Future studies that include more host attributes will also require larger sample sizes to account for the increased dimensionality.

## Material and methods

### Subject details

This single-center and non-interventional study was conducted in accordance with the Declaration of Helsinki Principles business ethics recommendations and Good Practices in Epidemiology. Written and informed consent was obtained from all participants before enrollment. Fifty-five (51) healthy Caucasian female volunteers living in Paris area participated in this observation. These women were enrolled according to age to obtain two groups of younger (20-26 yo, N=26) and older women (54-60 yo, N=25). Women in the older group were menopausal for at least one year. Non-inclusion criteria included pregnant or nursing women, women smoking or having stopped smoking for less than 5 years, women with facial cutaneous pathologies or pathologies affecting the skin, women under medication (such as corticosteroids, diuretics, antibiotics) during the last 2 years or for a treatment duration longer than 6 months, women under vitamin A treatment during the last 6 months or for a treatment duration longer than 1 month, women under medication (such as non-steroid anti-inflammatories, proton or anti-acid pump inhibitor) during the last 2 months or for a treatment duration longer than 8 days, women having applied on their arms a treatment (such as hormonal cream, AHA cream, antibiotics, antifungals) during the last 2 months or for a treatment duration longer than 8 days, women who have had UV sessions during the last 2 months or women with more than 48h-fever during the last 7 days.

### Measurement of biophysical parameters

Upon arrival at the laboratory with skin free of makeup and skincare products, subjects were acclimatized for 30 minutes before any measurements were performed under controlled environmental conditions with a temperature of 21 ± 2 °C and relative humidity at 50 ± 5 %. Skin capacitance (SC, determined with electric signal) was measured on the face using a corneometer (Dual MPA 580®, Monaderm). Collagen diffusion coefficient (CDC) and dermis water content (DWC) were determined on the face using an optical device (Dermo, Connected Physics). This device emits infra-red light and measures its absorption (relative to the dermis water content) and its scattering (relative to quality and quantity of collagen) into the skin.

### Sample collection and DNA extraction

Microbiome samples were collected on both cheeks using swabs and stored in preservative buffer at −80C until extraction. Extracted was performed using the GenElute Bacterial Genomic DNA kit (Sigma Aldrich) with the following modifications as previously published^37^: Swabs were placed in tubes containing 350 ul Tissue and Cell Lysis solution (Lucigen), 1.5 mg/ml lysozyme (Sigma Aldrich), 5 units of lysostaphin (Sigma Aldrich), 5 units of mutanolysin (Sigma Aldrich), and 100 ul of 0.1mM glass beads (Biospec). Swabs were incubated at 37 degrees Celsius for 30 minutes then bead beaten for 6 minutes at 30Hz in a TissueLyser II (Qiagen). The rest of the protocol was followed increasing the volumes to account for the additional lysis buffer. Tubes were spun down (1 minute at 15000 x g) in a microfuge prior to loading the columns to avoid any transfer of beads.

### mWGS sequencing and preprocessing

Sequencing libraries were prepared using the Nextera FLEX kit and Nextera Unique Dual Indexes (Illumina). Pooled libraries were sequenced on a NovaSeq 6000 (Illumina).

Demultiplexed Illumina reads were trimmed and quality checked with Cutadapt^38^ (v0.4.1), requiring a minimum length of 50 base pairs and a default Phred quality score of 20. Human reads were then removed with Bowtie2^39^ (v2.2.9, --very-sensitive mode), mapping to the hg19 human reference genome.

### Species-level community profiling

Community compositions were profiled using MetaPhlAn3.0^40^. To make sure that each subject was represented in the dataset only once, the community composition profiles of the left and right cheeks of the same subject were averaged for all analyses except when assessing intra-personal heterogeneity.

### Strain-level composition profiling

To estimate strain diversity, we used a reference-based method as described previously^7^. Reference databases were generated by compiling all Refseq genomes for *C. acnes* and *S. epidermidis*. Information for these genomes was available at https://github.com/ohlab/Strain_collection. Phylogenic trees were generated using Parsnp^41–44^ (v1.2, default parameters), and visualized with iToL^45^ (v3) (Supplementary Figure S3). Genomes were assigned into clades based on their primary branch from an unrooted tree.

Reads were then mapped to the genome databases for each species with Bowtie2^39^ (v2.3.4.3) using k=10 and “very-sensitive” mode. We then used Pathoscope^46^ (v2.0.6) on the resulting SAM files using default parameters for reassignment to nearest neighbor strains.

Determination of *C. acnes* phylotype and multi-locus sequence typing was conducted using PubMLST^47^ using two genomes representing the phylogenetic clade (Genbank assembly accession: GCA_002831705 and GCA_001750525).

### Microbial functional profiling

To characterize the general metabolic and functional composition of our dataset, reads were classified using HUMAnN2^48^ (v2.8.0, diamond mode). Profiling of ARG was conducted as described in Larson et al^7^. First, mWGS reads from all samples were pooled and assembled *de novo* using MEGAHIT^49,50^ (v1.0.6) with default parameters. Genes were then predicted from the resulting contigs using prodigal^51^ (v2.6.3) [53] under the “-meta” model with default parameters. The predicted genes were clustered using UCLUST^52^ (the cluster_fast algorithm in USEARCH v8.0.1517) at 90% sequence identity. Centroids were extracted from each cluster, from which antimicrobial resistance genes were annotated using DeepARG^53^ (v1.0.1, align mode for genes). Reads were mapped to the predicted AR genes using Bowtie2^39^ (v2.3.4.3, --very-sensitive mode) and counted using SAMtools^54^ (v1.10).

### Statistical analysis

All statistical analyses were performed in R^55^ (4.1.3) unless otherwise noted. False discovery rates were controlled using the Benjamini-Hochberg procedure. We reported results with p (or q) < 0.1 for univariate tests, and q < 0.25 for multivariate analyses (the default threshold adopted by MaAsLin2^22^). Differences in numerical variables observed between age groups (that is, not adjusted for other host attributes) were tested using the Wilcoxon Rank Sum test (the “wilcox.test” function in R). Correlations between host attributes were tested using the “cor” function in R. The variance-inflation factors were calculated using the “vif” function in the R package “car”^56^ (v3.0.12). Univariate multiple regression models were fit using the “lm” function in R, with type II ANOVA tests performed using the “Anova” function in the R package “lmerTest”^57^ (v3.1.3). When the dependent variable is the relative abundance of a microbial taxon, the relative abundance values were log-transformed. Multivariate multiple regression models were fit using MaAsLin2^22^ (v1.8.0), with default parameters. Shannon’s index and Bray-Curtis dissimilarity were estimated using the R package “vegan”^58^ (v2.5.7). Non-metric multidimensional scaling (NMDS) was performed using the metaMDS function in the R package “vegan”^58^ (v2.5.7). dbRDA was also performed using the R package “vegan”^58^ (v2.5.7) based on Bray-Curtis dissimilarity.

## Supplementary material

Supplementary Table S1. Host attributes and their variance inflation factors (VIF).

Supplementary Table S2. Microbial read counts of the metagenomic shotgun samples.

Supplementary Figure S3. Phylogenetic clades of *C. acnes* and *S. epidermidis*. Reference genomes used in this study were assigned into clades based on their primary branch from an unrooted tree. This figure was adapted from Figure S15 of Larson et al^7^.

## Notes

### Competing Interest Statement

The authors have declared no competing interest.

