## Supplementary figures and images for "Skin microbiome attributes associate with biophysical skin aging"

### Supplementary Figure S3

*C. acnes*

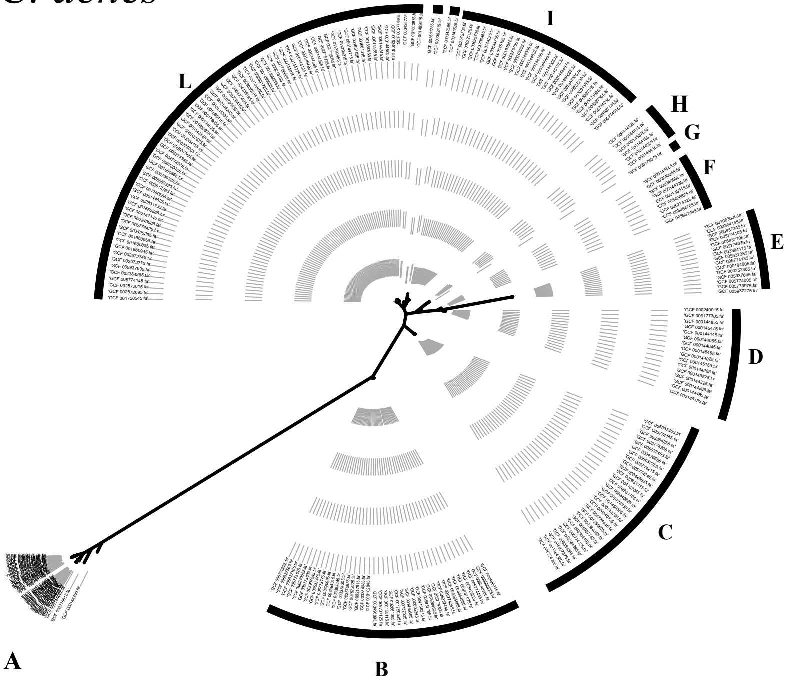


*S. epidermidis*

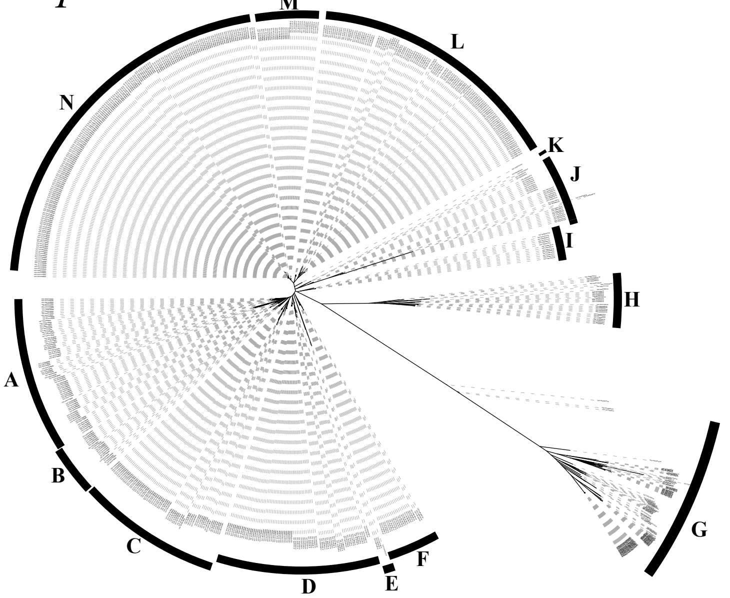
